# A bacteria-based assay to study SARS-CoV-2 protein-protein interactions

**DOI:** 10.1101/2021.10.07.463611

**Authors:** Benjamin L. Springstein, Padraig Deighan, Grzegorz Grabe, Ann Hochschild

## Abstract

Methods for detecting and dissecting the interactions of virally encoded proteins are essential for probing basic viral biology and providing a foundation for therapeutic advances. The dearth of targeted therapeutics for the treatment of COVID-19, an ongoing global health crisis, underscores the importance of gaining a deeper understanding of the interactions of SARS-CoV-2-encoded proteins. Here we describe the use of a convenient bacteria-based two-hybrid (B2H) system to analyze the SARS-CoV-2 proteome. We identify sixteen distinct intraviral protein-protein interactions (PPIs), involving sixteen proteins. We find that many of the identified proteins interact with more than one partner. We further show how our system facilitates the genetic dissection of these interactions, enabling the identification of selectively disruptive mutations. We also describe a modified B2H system that permits the detection of disulfide bond-dependent PPIs in the normally reducing *Escherichia coli* cytoplasm and we use this system to detect the interaction of the SARS-CoV-2 spike protein receptor-binding domain (RBD) with its cognate cell surface receptor ACE2. We then examine how the RBD-ACE2 interaction is perturbed by several RBD amino acid substitutions found in currently circulating SARS-CoV-2 variants. Our findings illustrate the utility of a genetically tractable bacterial system for probing the interactions of viral proteins and investigating the effects of emerging mutations. In principle, the system could also facilitate the identification of potential therapeutics that disrupt specific interactions of virally encoded proteins. More generally, our findings establish the feasibility of using a B2H system to detect and dissect disulfide bond-dependent interactions of eukaryotic proteins.

**Importance:** Understanding how virally encoded proteins interact with one another is essential in elucidating basic viral biology, providing a foundation for therapeutic discovery. Here we describe the use of a versatile bacteria-based system to investigate the interactions of the protein set encoded by SARS-CoV-2, the virus responsible for the current pandemic. We identify sixteen distinct intraviral protein-protein interactions, involving sixteen proteins, many of which interact with more than one partner. Our system facilitates the genetic dissection of these interactions, enabling the identification of selectively disruptive mutations. We also describe a modified version of our bacteria-based system that permits detection of the interaction between the SARS-CoV-2 spike protein (specifically its receptor binding domain) and its cognate human cell surface receptor ACE2 and we investigate the effects of spike mutations found in currently circulating SARS-CoV-2 variants. Our findings illustrate the general utility of our system for probing the interactions of virally encoded proteins.

## Introduction

The causative agent of COVID-19, SARS-CoV-2, like SARS-CoV (hereafter SARS-CoV-1) and the Middle East respiratory syndrome coronavirus (MERS-CoV), is a zoonotic pathogen that belongs to the genus of β-coronaviruses [1,2]. A ~30 kb single stranded (+)-sense RNA virus, SARS-CoV-2 encodes 16 non-structural proteins (Nsp1-Nsp16), which are transcribed from two major open reading frames (ORF1a and ORF1b) and later post-translationally processed by proteases to give rise to the individual Nsps [3]. The main function of the Nsps is to provide and maintain the replication and transcription complex (RTC), promoting viral RNA synthesis by the RNA-dependent RNA polymerase Nsp12 [3]. However, the Nsps have also been implicated in other viral processes such as host innate immune system evasion – for example, by suppressing aspects of the interferon response [4]. The virus also encodes four structural proteins, the membrane (M) protein, the nucleocapsid (N) protein, the envelope (E) protein and the spike (S) glycoprotein, and at least six accessory proteins (ORF3a, ORF6, ORF7a, ORF7b, ORF8 and ORF10) [5]. The main function of coronavirus structural proteins is to mediate cell entry, virus particle assembly and release from the host cells by budding, though like the Nsps, structural proteins also participate in immune evasion. By contrast, the accessory proteins are non-conserved and highly variable among different coronavirus species; although their functional roles remain largely unknown, they too have been associated with immune evasion and disease severity [3].

Given the ongoing global crisis caused by the SARS-CoV-2 pandemic and the continuing need for targeted therapeutics for the treatment of COVID-19, understanding the intraviral and viral-host protein-protein interactions (PPIs) of SARS-CoV-2 remains a priority. An extensive virus-virus and host-virus PPI study recently highlighted the importance of Nsp10 as a potential inducer of the so-called cytokine storm (a dysregulated and hyperactive immune response) [6], thought to be the main cause for severe disease outcome and death in COVID-19 patients [2]. Li *et al.* (2021) further identified Nsp8 as a SARS-CoV-2 PPI hub [6], promoting interactions with other Nsps, accessory proteins, and one structural protein. Similar observations were previously also obtained for SARS-CoV-1 Nsp8 [7]. These findings suggest that Nsp8 and Nsp10 might provide particularly efficacious targets for drug development.

The SARS-CoV-2 spike protein, which is present on the viral surface as trimers, consists of two functionally distinct subunits, S1 and S2 [8,9]. The membrane-distal S1 subunit uses its receptor-binding domain (RBD) to initiate the process of viral entry into human host cells by binding to the cell surface protein angiotensin-converting enzyme 2 (ACE2), which also serves as the receptor for SARS-CoV-1 but not for the more distantly related MERS-CoV. Following ACE2 binding, the membrane-localized host cell serine protease TMPRSS2 cleaves the spike protein at a specific site, triggering a series of dramatic conformational changes in the S2 subunit, which in turn mediate fusion of the viral and host membranes, enabling viral entry [10]. As well as being a critical determinant of viral tropism, the RBD is a major target for SARS-CoV-2-neutralizing antibodies, including those identified from convalescent patient peripheral blood mononuclear cells and those elicited by current (spike-based) vaccines [9,11–18].

Compared with those of other RNA viruses, the mutation rate of SARS-CoV-2 is considered low-moderate (6-9×10^-4^ bases/genome/year) [19–21], although others have pointed out that multiple identical mutation hotspot events occurring at different points in time could lead to an underestimation of the overall mutation rate [22]. Nevertheless, the pandemic has given rise to a proliferation of variant lineages, including those designated variants of concern (VOCs) by the World Health Organization (WHO), based on one or more of the following criteria: increase in transmissibility; increase in virulence; decrease in effectiveness of public health measures, diagnostics, therapeutics or vaccines (www.who.int/en/activities/tracking-SARS-CoV-2-variants). All of the VOCs carry spike mutations, including one or more that localize to the RBD, motivating efforts to gain a systematic understanding of the effects of RBD amino acid substitutions on ACE2 binding [23].

Here we employ a bacterial two-hybrid (B2H) system [24,25] to study the PPIs of SARS-CoV-2 in a heterologous non-eukaryotic system. Using this system, we describe a bacteria-based intraviral interactome. We further demonstrate the utility of the bacterial system for genetically dissecting the SARS-CoV-2 PPIs by identifying mutations that selectively affect one or another interaction. In addition, we describe a modified B2H system that allows us to detect disulfide bond-dependent PPIs in the otherwise reducing *Escherichia coli* cytoplasm. We use this system to detect the spike RBD-ACE2 interaction and to investigate the effects of mutations found in VOCs. Our findings set the stage for further investigations of viral PPIs in a convenient and genetically tractable bacterial system, as well as establishing the feasibility of using our modified system to detect and dissect disulfide bond-dependent PPIs of other eukaryotic proteins.

## Results

### Bacterial two-hybrid system to detect interactions of SARS-CoV-2 proteome

Previous studies have used yeast two-hybrid (Y2H) systems, a mammalian two-hybrid system and co-immunoprecipitation experiments (co-IPs) to investigate the SARS-CoV-1 and SARS-CoV-2 protein interactomes, identifying overlapping but also distinct interactions depending on the employed system [6,7,26,27]. Compared with bacteria, yeast have a relatively slow growth rate and are more difficult to culture and transform for labs that do not routinely work with yeast. To provide a more accessible alternative to Y2H systems as well as the less commonly used mammalian two-hybrid system, we here describe the successful use of a B2H system developed in our lab (Fig. 1A) [24,25] to test for viral PPIs. We fused all NCBI-predicted *E. coli* codon-optimized SARS-CoV-2 open reading frames (ORFs; listed in Fig. 1B, see also NCBI accession #: NC_045512.2) to the DNA binding protein CI of bacteriophage λ (λCI) and to the N-terminal domain of the α subunit (αNTD) of RNA polymerase (RNAP). We then tested each SARS-CoV-2 ORF for interaction with the other SARS-CoV-2 ORFs and itself. Interaction between two given ORFs (X and Y), fused to αNTD and λCI, respectively, stabilizes the binding of RNAP to the test promoter such that the magnitude of the *lacZ* reporter gene expression correlates with the strength of the PPI (Fig. 1A).

**Fig. 1:**
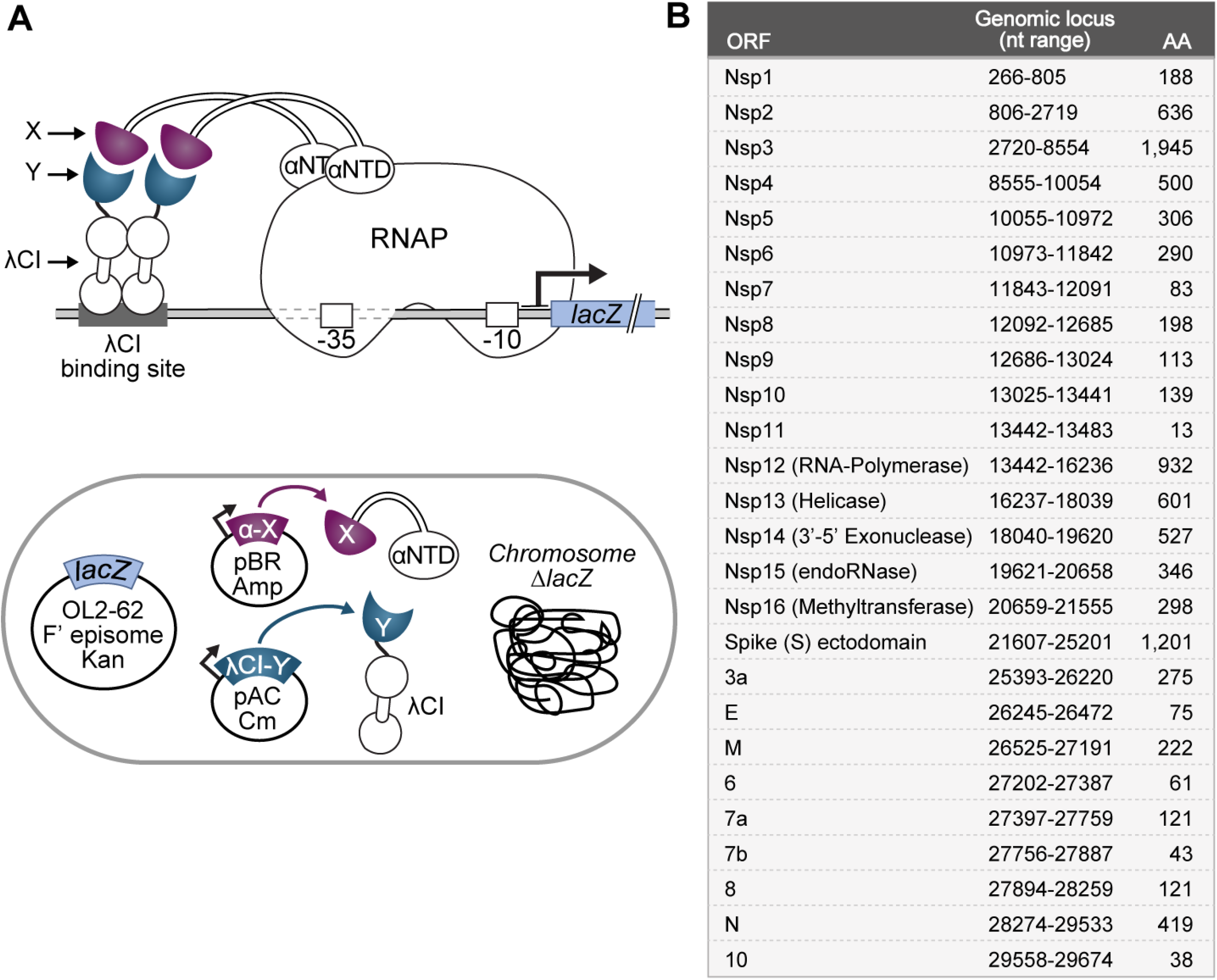
Bacterial two-hybrid assay used to study the SARS-CoV-2 interactome. (**A**) (top) Schematic depiction of the employed transcription-based bacterial two-hybrid system. Interaction between protein moieties X (purple) and Y (slate blue), which are fused to the N-terminal domain of the *α* subunit of *E. coli* RNAP (αNTD) and the λCI protein, respectively, stabilizes the binding of RNAP to test promoter p*lac*O_L_2-62, thereby activating transcription of the *lacZ* reporter gene. The test promoter bears the λ operator OL2 centered at position −62 upstream of the transcription start site. (bottom) *E. coli* cell containing genetic elements that are involved in the bacterial two-hybrid system. The chromosomal *lacZ* locus is deleted and the test promoter and fused *lacZ* reporter gene are encoded on an F’ episome. The λCI-Y and αNTD-X fusion proteins are encoded on compatible plasmids and produced under the control of IPTG-inducible promoters. (**B**) List of all tested SARS-CoV-2 ORFs as predicted by the NCBI reference genome (Accession #: NC_045512.2). The respective nucleotide range for each ORF based on the NCBI reference sequence is indicated, together with the resulting amino acid sequence length. Except for the spike protein, all ORFs were cloned as full-length genes. For spike, we chose to test the interaction of its ectodomain (aa 16-1213) to avoid complications due to its N-terminal signal peptide and C-terminal transmembrane domain.

### Identification of the SARS-CoV-2 interactome using a B2H system

Using our B2H system, we initially tested each SARS-CoV-2 ORF against each other SARS-CoV-2 ORF and itself in biological duplicate. Protein pairs with at least a 2-fold activation of *lacZ* over background in one of the replicates were selected for further analysis. The list of interacting proteins was further refined by performing repeat experiments with three biological replicates for each initially identified potential PPI pair. This resulted in a final list of sixteen interacting SARS-CoV-2 protein pairs, including four self-interactions (Fig. 2). Some of these interactions were identified only with a specific fusion partner combination *(i.e.,* protein X fused to αNTD and protein Y fused to λCI, or the other way around), while others were fusion partner-independent *(i.e.,* interaction between proteins X and Y regardless of their fusion to αNTD or λCI). Self-interacting proteins (Nsp7, Nsp9, ORF6 and ORF10) were by definition fusion partner-insensitive; however, four other pairs of proteins (Nsp7+Nsp8, Nsp10+Nsp14, Nsp10+Nsp16, Nsp3+N, and Nsp8+ORF6) also interacted detectably regardless of the fusion partner (Supplementary Fig. 1).

**Fig. 2:**
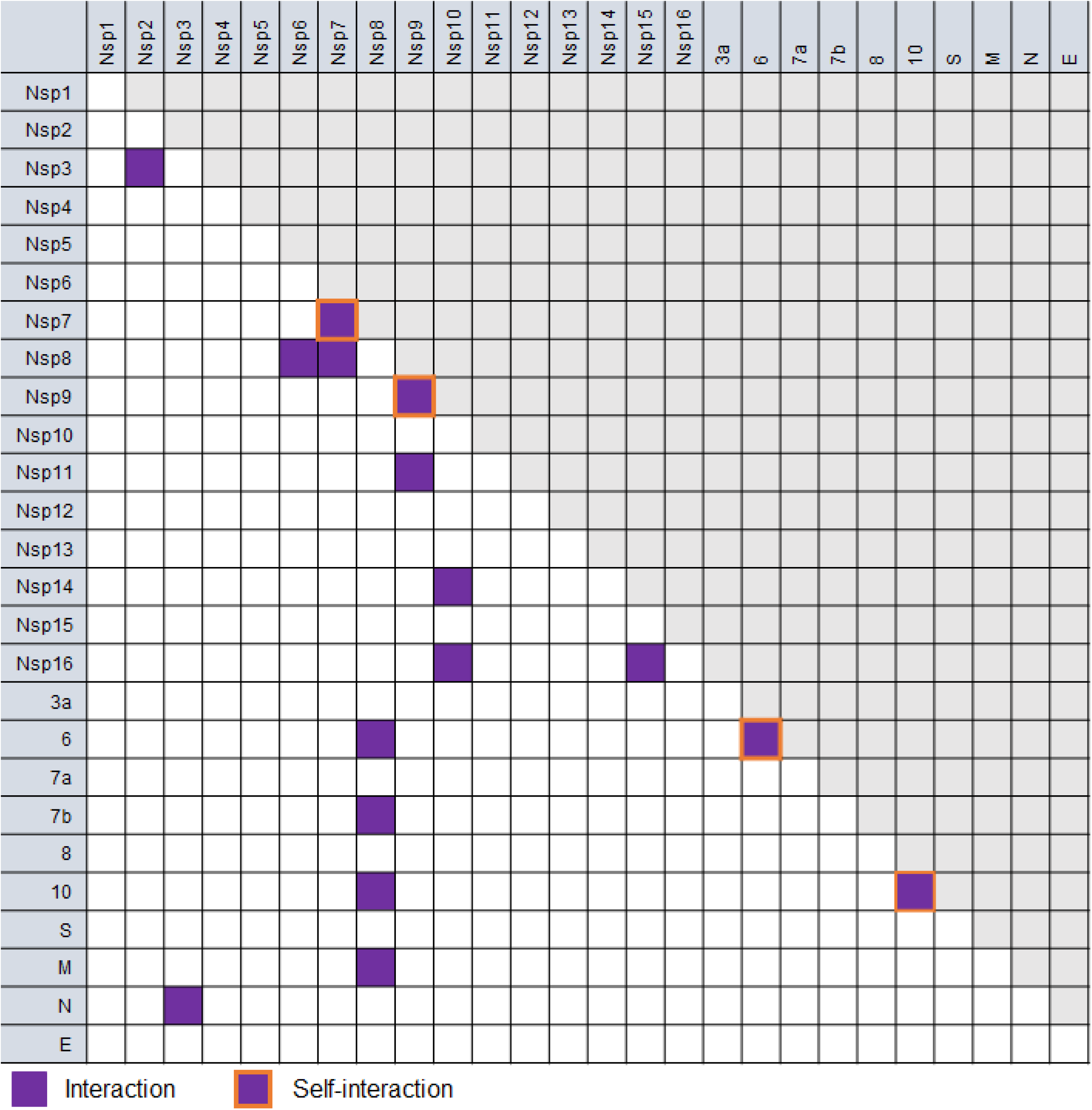
Detection of protein-protein interactions by the bacterial two-hybrid system. Interaction matrix of all tested ORFs. Positive interactions, regardless of the fusion partner, are indicated with purple squares and self-interactions are indicated by orange-framed squares. Detailed information about fusion constructs for which positive interactions were identified is given in Supplementary Fig. 1. To avoid data duplication, only one half of the matrix is shown while the other is shaded in grey.

Among the identified interacting pairs, several particularly strong PPIs were observed, including the Nsp7 self-interaction, Nsp7+Nsp8, Nsp10+Nsp16, N+Nsp3 and Nsp9+Nsp11 (Fig. 3). In fact, the Nsp10+Nsp16 pair interacted significantly more strongly than our positive control, representing one of the strongest interactions we have ever measured with our B2H assay. For our B2H assays, we routinely consider an interaction to be reliable when we detect at least a two-fold increase in *lacZ* reporter gene expression (measured as β-galactosidase activity) over the background (obtained with the negative controls). Applying this cut-off to our experimental data, we identified several medium-to-weak interactions (2-to 5-fold increase over the negative controls; Supplementary Fig. 2). The interactions of Nsp8+ORF7b and ORF10+ORF10 closely missed the 2-fold cutoff but were nonetheless included in the list because a previous SARS-CoV-2 interactome study also identified those interactions (based on co-IP data) [6].

**Fig. 3:**
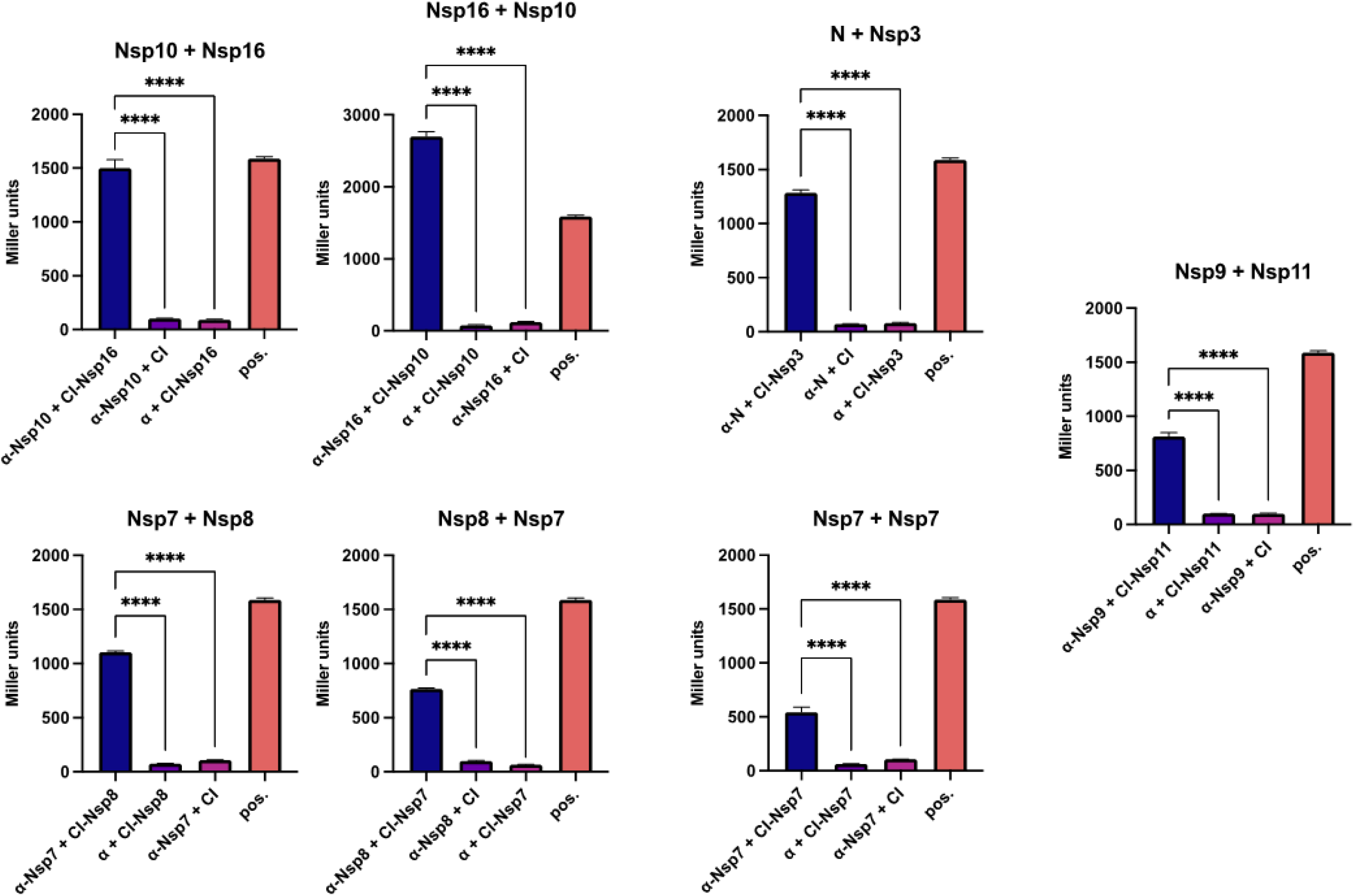
Strong SARS-CoV-2 protein-protein interactions identified by B2H assays. Shown are two-hybrid data for strong interactions (arbitrarily defined as Miller units > 500). Indicated ORFs are fused either to the αNTD (indicated as α) or to full-length λCI (indicated as CI). α and λCI negative controls express full-length a and full-length λCI, respectively. The interaction of domain 4 of the RNAP σ^70^ subunit (fused to the αNTD) with the flap domain of the RNAP β subunit (fused to λCI) served as a positive control (pos) [81,82]. Bar graphs show the averages of three biological replicates (n=3) and β-galactosidase activities are given in Miller units. Error bars indicate the standard deviation. Values indicated with asterisks are significantly different from the negative control. ****: P<0.0001 (One-way ANOVA with Turkey’s multiple comparison test).

Comparison of our SARS-CoV-2 B2H data with the previously reported SARS-CoV-2 Y2H and co-IP data [6] revealed four PPIs that were shared among the three assay systems, providing strong support for their biological relevance (Supplementary Fig. 3). These included Nsp7+Nsp8, Nsp8+ORF10, Nsp10+Nsp14 and ORF6+ORF6. Others were identified either in only one of the assay systems *(i.e.,* B2H, Y2H or co-IP) or in two assay systems (B2H and Y2H, B2H and co-IP, or Y2H and co-IP) (Supplementary Fig. 3). Furthermore, some of our identified interactions are validated by co-crystal structures. These included Nsp7+Nsp8 (Protein Data Bank (PDB) accession number 6YHU [28]), Nsp10+Nsp14 (PDB: 5NFY from SARS-CoV-1 [29] or more recently 7DIY from SARS-CoV-2 [30]), Nsp10+Nsp16 (PDB: 6W4H [31]) and the Nsp9 self-interaction (PDB: 6W9Q [32]). Notably, no self-interaction of Nsp9 was identified in a previous Y2H and co-IP analysis of the SARS-CoV-2 interactome [6], highlighting the importance of employing several different interaction assays when studying the interactome of a given protein set to avoid loss of information due to experimental system idiosyncrasies.

Similar to previous observations for SARS-CoV-1 [7], we identified Nsp8 as a major SARS-CoV-2 interaction hub, interacting with six other SARS-CoV-2 ORFs (Fig. 3, Supplementary Fig. 2), consistent with a critical role for Nsp8 in SARS coronavirus biology. Nonetheless, most of the interaction partners we identified for Nsp8 in SARS-CoV-2 are different than those identified previously for SARS-CoV-1 [7,26,27] (Supplementary Fig. 4). Overall, only six PPIs were identified in our SARS-CoV-2 B2H analysis and at least one of three independent SARS-CoV-1 Y2H studies, including two involving Nsp8 (Supplementary Fig. 4). Notably, there are considerable differences between the results of the three previous Y2H studies [7,26,27] and only three PPIs (Nsp8+Nsp7, Nsp10+Nsp14, and Nsp10+Nsp16) were independently identified in two SARS-CoV-1 two-hybrid assays and our SARS-CoV-2 B2H assay (Supplementary Fig. 4). This could reflect significant differences between the PPI networks in SARS-CoV-1 and SARS-CoV-2 and/or differences in the assays themselves (procedures and background organism).

### Targeted mutational screens identify interaction partner-specific sites of protein-protein interaction in CoV-2 proteins with more than one interaction partner

As a genetic assay, the B2H system facilitates the dissection of specific PPIs through both targeted and random mutagenesis. Having established the utility of the B2H assay in testing for viral PPIs, we next sought to use this assay to dissect the interactions of selected viral proteins through targeted mutational analysis. Specifically, we chose proteins that interacted with more than one partner and sought to disrupt the interaction of such a protein with one of its partners while preserving its interaction with another. We initially selected Nsp10 with two known interaction partners, Nsp14 and Nsp16, and attempted to disrupt only its interaction with Nsp14. To identify suitable targets for mutagenesis, we analyzed the crystal structures of Nsp10-Nsp14 (PDB ID: 5NFY [29]) and Nsp10-Nsp16 (PDB ID: 6W4H [31]) and their protein-protein interfaces using PDBePISA [33]. Based on this approach, we selected three sets of amino acid substitutions likely to affect the binding of Nsp10 to Nsp14, while leaving its interaction with Nsp16 intact (assuming that the substitutions do not result in allosteric effects). While the Nsp10 F16A/F19A/V21A set targeted the hydrophobic region, the Nsp10 T5A/T12A/S15A and S29A/S33A sets partially disrupted the hydrogen bond network of the Nsp10-Nsp14 interface (Fig. 4A,C). Each of the three multiply substituted Nsp10 mutants lost the ability to interact detectably with Nsp14 while maintaining an approximately wild-type interaction with Nsp16 (Fig. 4B). We note that the close approach of amino acid side chains at a protein-protein interface as revealed by X-ray crystallography does not necessarily indicate that they participate in a functionally important interaction. However, the loss of a detectable interaction between each of the three Nsp10 mutants and Nsp14 in our B2H assay suggests that at least a subset of the selected residues make stabilizing contacts. Furthermore, although the Nsp10-Nsp16 interaction serves as a control, we also confirmed that the introduced amino acid substitutions were not generally destabilizing (Supplementary Fig. 5).

**Fig. 4:**
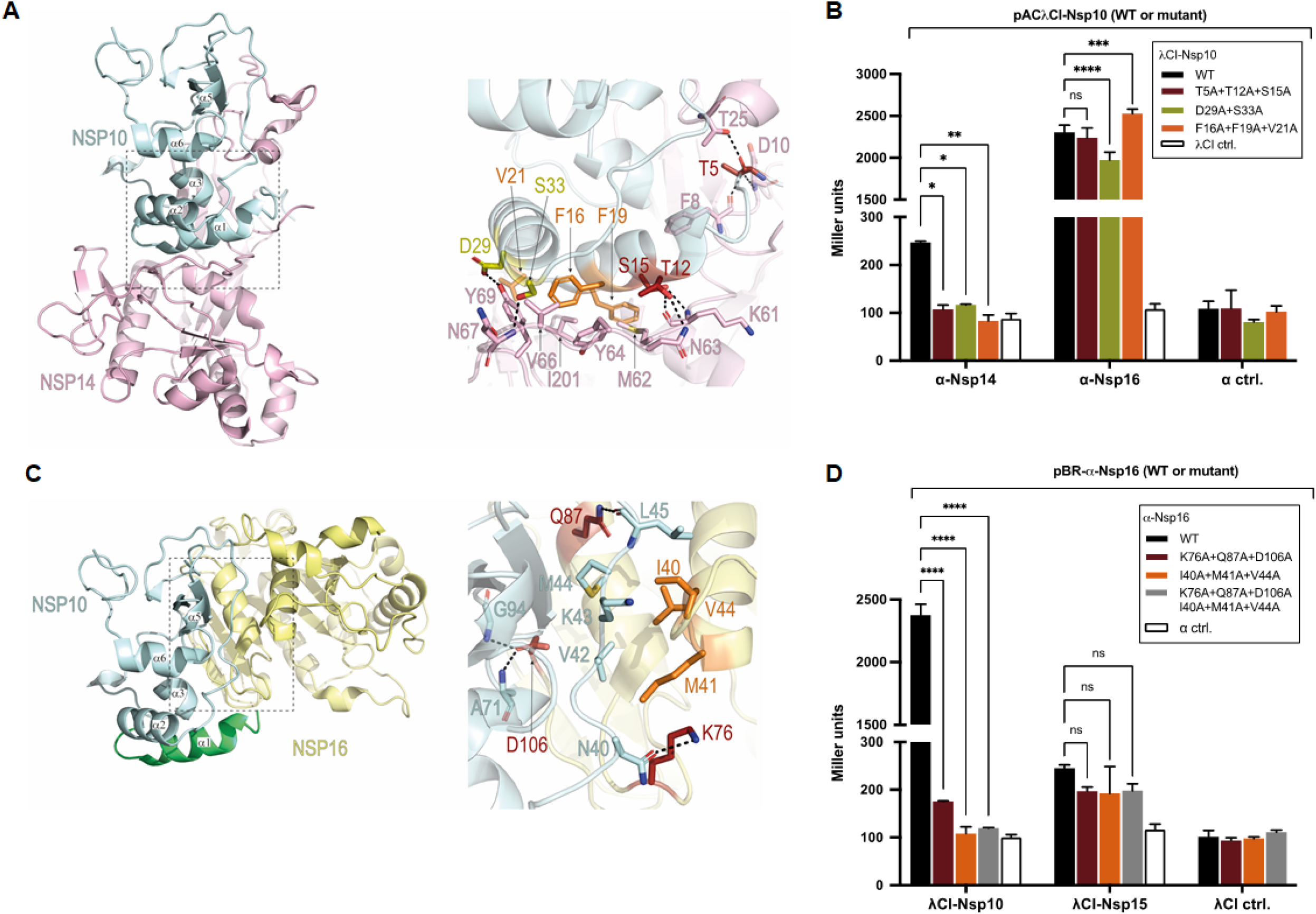
Selective disruption of protein interfaces for protein with two interaction partners. (**A**) Depiction of crystal structure (PDB ID: 5NFY [29]) of SARS-CoV-1 Nsp10 (pale cyan) in complex with Nsp14 (pale pink). Zoom-in shows amino acids (sticks) chosen for mutational analysis of Nsp10 (orange, olive, and burgundy) and their corresponding main interaction partners in Nsp14 (pale pink). (**B**) B2H results showing effects of Nsp10 substitutions on its interactions with Nsp14 and with Nsp16. Amino acid substitutions introduced into Nsp10 are given in the box. (**C**) Depiction of crystal structure of the SARS-CoV-2 Nsp16-Nsp10 protein complex (PDB ID: 6W4H [31]) colored respectively in pale yellow and pale cyan. Additional N-terminal Nsp10^7-22^ region is included and was obtained from superimposed Nsp10 structure from PDB ID: 5NFY (green). Zoom-in shows amino acids (sticks) chosen for mutational analysis of Nsp16 (orange and burgundy) and their corresponding main interaction partners in Nsp10 (pale cyan). (**D**) B2H results showing effects of Nsp16 substitutions on its interactions with Nsp10 and with Nsp15. Amino acid substitutions introduced into Nsp16 are given in the box. (**B,D**) Indicated ORFs are fused either to the αNTD (indicated as α) or to full-length λCI. α and λCI negative controls express full-length α and full length λCI, respectively. Bar graphs show the averages of three biological replicates (n=3) and β-galactosidase activities are given in Miller units. Error bars indicate the standard deviation. Values indicated with asterisks are significantly different from the WT. ns: not significant; *: P<0.05; **: P<0.01; ****: P<0.0001 (One-way ANOVA with Dunnett’s multiple comparison test). Black dashed lines in **A** and **C** represent hydrogen bonds.

We then focused on Nsp16 with two interaction partners, Nsp10 and Nsp15, targeting the Nsp16-Nsp10 pair, which displayed a significantly higher B2H signal than that of the Nsp16-Nsp15 pair. Here we also utilized the available crystal structure for the Nsp16-Nsp10 complex; however, as there is no structure for the Nsp16-Nsp15 complex, the substitutions introduced into Nsp16 were based solely on their predicted effects on its interaction with Nsp10. Endeavoring to disrupt the Nsp16-Nsp10 interaction, we created two Nsp16 triple substitution mutants, targeting hydrophobic (I40A/M41A/V44A) or hydrophilic (K76A/Q87A/D106A) contacts, and a mutant with the six substitutions combined (Fig. 4C). The data reveal drastic effects of these substitutions on the binding of Nsp16 to Nsp10, resulting in near background or background levels of reporter gene expression for each of the mutants (Fig. 4D). The effects of the same substitutions on the binding of Nsp16 to Nsp15 were modest and not statistically significant. Notably, even though Nsp16 interacts much more weakly with Nsp15 than with Nsp10, reporter gene expression was lower for each of the Nsp16 mutants in combination with Nsp10 than when tested in combination with Nsp15 (Fig. 4D). We also confirmed that these effects are not the result of altered protein levels (Supplementary Fig. 6). Together, these data illustrate a proof-of-principle approach that can be used to obtain functionally informative mutants within a PPI network.

### The B2H system as a tool to study circulating spike variants and their binding to ACE2

To further assess whether our B2H system can facilitate the study of emerging mutational changes in viral populations, we next asked whether we could use our system to study the interaction between the SARS-CoV-2 spike protein and ACE2. For this, we obtained an *E. coli* codon-optimized gene fragment encoding the human ACE2 peptidase domain (aa 19-615, hereafter ACE2). We inserted this gene fragment and a set of gene fragments encoding multiple domains of the spike protein (including the RBD, aa 331-521) into our two-hybrid vectors, fusing ACE2 and each of the spike domains to both λCI and αNTD. Initial experiments using our standard *E. coli* B2H strain (FW102 OL2-62 termed B2H; Supplementary Table 1A) failed to reveal an interaction of ACE2 with any of the selected spike domains (Supplementary Fig. 7; data not shown). However, previous studies demonstrated that proper disulfide bond formation is essential in order for spike and ACE2 to engage in a direct interaction [34,35]. Because the *E. coli* cytoplasm is a reducing environment [36], we considered the possibility that the failure to detect a spike-ACE2 interaction with our standard B2H strain might be due to a lack of proper disulfide bond formation. To circumvent this obstacle, we modified a commercially available *E. coli* strain (SHuffle from NEB, MA, USA) that permits the efficient expression and formation of active full-length antibodies in *the E. coli* cytoplasm [37], adapting it for use with our two-hybrid system (Methods). The SHuffle strain is deleted for two genes that encode cytoplasmic reductases *(trxB* and *gor)* and also harbors the normally periplasmic disulfide bond isomerase DsbC in the cytoplasm [38,39]. With this modified oxidizing strain (termed BLS148; Supplementary Table 1A), we were able to detect an interaction of the spike RBD with ACE2 (Fig. 5A,B). Moreover, this interaction was abrogated when we mutated a pair of cysteine residues (replacing them individually and in combination with serine residues) that engage in disulfide bond formation within the RBD (C379 and C432) [34], consistent with the surmise that the oxidizing strain permits detection of the RBD-ACE2 interaction by enabling appropriate disulfide bond formation and correct folding of the interacting partners (Fig. 5B; Supplementary Fig. 8).

**Fig. 5:**
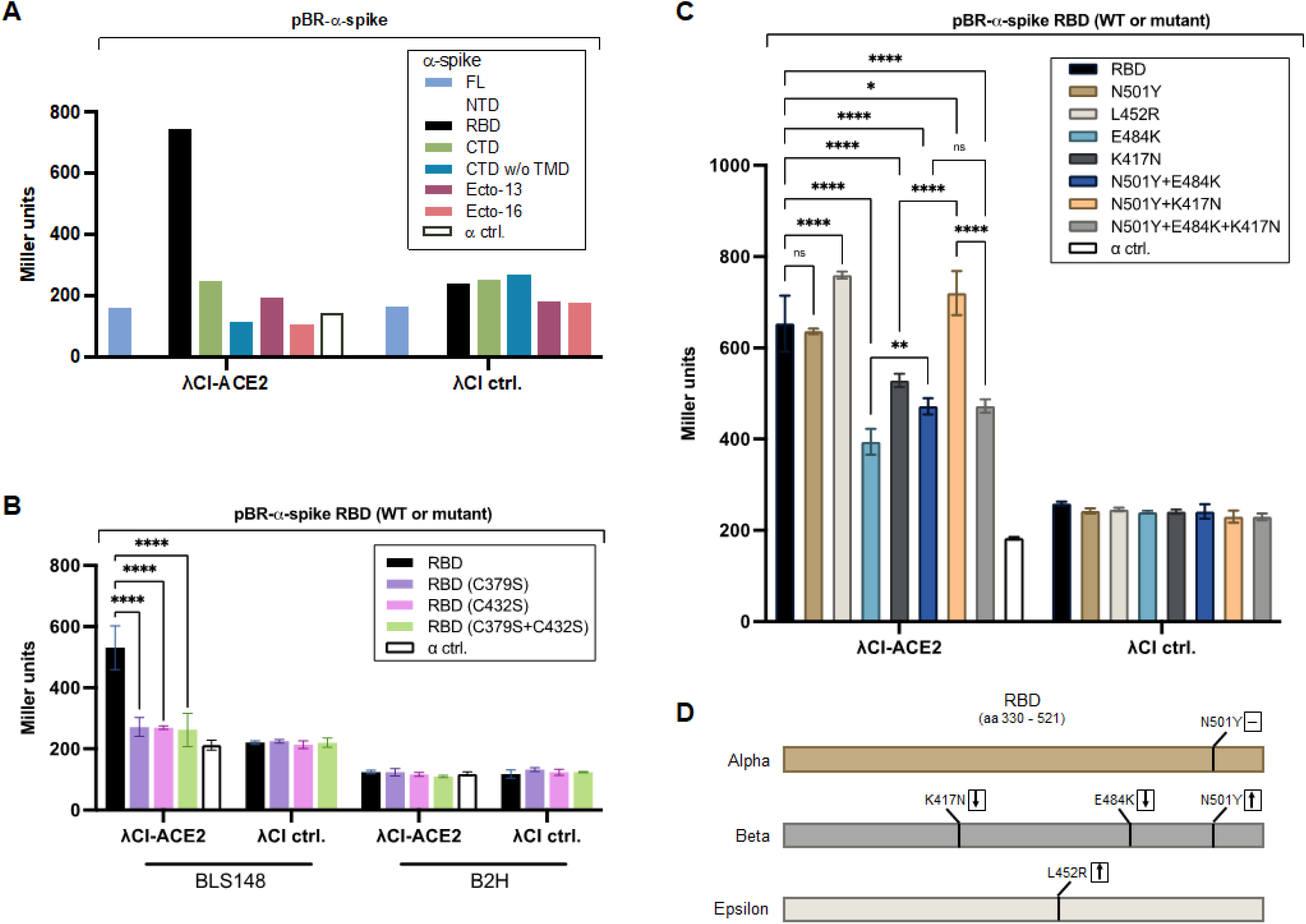
Interaction of spike RBD and ACE2 in an oxidizing *E. coli* strain. (**A**) Bacterial two-hybrid assays of (**A**) spike domains (as listed in Supplementary Fig. 7) tested against ACE2 in BLS148, (**B**) indicated spike RBD cysteine mutants tested against ACE2 in BLS148 and B2H or (**C**) indicated spike RBD circulating variants tested against ACE2 in BLS148. FL: full-length, NTD: N-terminal domain; RBD: receptor binding domain; CTD: C-terminal domain (with or without transmembrane domain (TMD)), Ecto: Ectodomain starting either at aa 13 or 16. (D) Schematic depicting amino acid substitutions present in each of three RBD variants tested. The measured effect of each substitution on ACE2 binding is indicated with a dash (no effect), a downward pointing arrow (weakened binding) or an upward pointing arrow (strengthened binding). (**A-C**) spike domains or RBD mutant variants were fused to the αNTD (indicated as α) and ACE2 was fused to full-length λCI. α and λCI negative controls express full-length α and full-length λCI, respectively. Bar graphs show (**A**) one biological replicate or (**B,C**) the averages of three biological replicates (n=3) and β-galactosidase activities are given in Miller units. Note: results depicted in (**C**) have been confirmed in a total of seven independent experiments, one of which is shown here. Error bars indicate the standard deviation. Values indicated with asterisks are significantly different from the negative control. ns: not significant; *: P<0.05; **: P<0.01; ****: P<0.0001 (Two-way ANOVA with Turkey’s multiple comparison test). Western blot analysis indicated that the spike RBD mutants used in (**B**) and (**C**) are present at intracellular levels comparable to the wild-type RBD, ruling out protein instability as a cause for the observed effects (Supplementary Figs. 8 and 9).

Having adapted our B2H system for the study of disulfide bond-dependent PPIs, we sought to test different spike (RBD) circulating variants for their abilities to bind ACE2. The RBD amino acid substitutions included in our study are found in several SARS-CoV-2 variants that were previously designated VOCs by the Centers for Disease Control and Prevention (USA; https://www.cdc.gov/coronavirus/2019-ncov/variants/variant-info.html, initially accessed 05/30/2021). Specifically, we included the Alpha variant (B.1.1.7, first identified in the United Kingdom) that carries the RBD N501Y substitution, the Beta variant (B.1.351, first identified in South Africa) that carries the RBD K417N, E484K and N501Y substitutions, as well as the Epsilon variant (B.1.429, first identified in California) that carries the RBD L452R substitution, introducing the corresponding mutations into the spike RBD on our B2H vector (Fig. 5D). The latter two variants have recently gained more attention as they are considered immune escape variants, potentially resulting in a partial loss of immunity in previously infected or immunized people [40–47]. In contrast, the Alpha variant is not characterized by a marked escape from antibody neutralization [41,42,45–47]. Factors that are believed to contribute, potentially, to immune escape include changes in the spike protein that: (i) enhance or stabilize its binding to ACE2 or (ii) decrease the binding of specific anti-spike neutralizing antibodies [48–50].

As well as testing the Alpha, Beta and Epsilon RBDs for their abilities to bind ACE2, we included RBD mutants bearing component single and double substitutions from the Beta variant (Fig. 5C; Supplementary Fig. 9). We found that the N501Y substitution (in the context of the Alpha variant) had no observable effect on ACE2 binding. In contrast, the L452R substitution (Epsilon variant) resulted in a statistically significant increase in ACE2 binding. Substitutions K417N, E484K and N501Y (Beta variant) together resulted in a significant reduction in ACE2 binding, as did the individual component substitutions K417N and E484K (with the E484K substitution having the stronger effect). However, the effects of these two substitutions were partially (E484K) or fully (K417N) abrogated when combined with the N501Y substitution. The binding of the triply substituted variant was indistinguishable from that of the E484K/N501Y double mutant, indicating that in this context the K417N substitution neither weakens or strengthens the interaction. Together, these findings indicate that our modified B2H system enables detection of disulfide bond-dependent PPIs and can be used to investigate the effects of RBD variant substitutions on the RBD-ACE2 interaction.

## Discussion

### Use of bacteria-based assay to investigate SARS-CoV-2 interactome

Here we use a versatile bacteria-based genetic tool for detecting and dissecting PPIs [24,25] to screen the SARS-CoV-2 proteome for intraviral PPIs. We detected a total of sixteen PPIs, including four self-interactions. Nine of these interactions were also detected in a previous SARS-CoV-2 PPI study (Supplementary Fig. 3), as assessed by Y2H-based screens and/or mammalian cell-based co-IP experiments [6]. Additionally, four of the interactions we detected have been captured in co-crystal structures, including the Nsp9 self-interaction (PDB: 6W9Q; [32]), which was not identified by either Y2H or co-IP analyses [6]. Of the six interactions we detected that were not previously described in the context of SARS-CoV-2, three were previously detected by Y2H analyses in the context of SARS CoV-1 (Supplementary Fig. 4). Of the remaining three interactions, not previously described, two (Nsp9+Nsp11 and Nsp3+N) were particularly strong as assessed in our B2H assay (Fig. 3).

Although the different assays that have been used to characterize the SARS-CoV-2 interactome have provided results that often corroborate one another, there are many examples of interactions that have been detected with only one of the assays. These discrepancies highlight the importance of employing multiple assay systems, each with its own inherent limitations, to maximize the likelihood of obtaining a complete picture. Because of its experimental accessibility, we expect that our B2H assay will be useful in evaluating other viral proteomes, particularly by taking advantage of our oxidizing reporter strain that better approximates the eukaryotic cell environment in allowing for proper disulfide bond formation in the *E. coli* cytoplasm [51,52]. To further extend the spectrum of testable viral and eukaryotic PPIs, the system could be augmented to enable the detection of phosphorylation-dependent PPIs by introducing specific mammalian kinases into our reporter stain [53]. Given that many mammalian (and presumably viral) proteins are constitutively phosphorylated in yeast [54–56], a lack of properly phosphorylated proteins in our B2H system could explain, at least in principle, why some SARS-CoV-2 PPIs were identified only in the Y2H screens [6] and not in our system (Supplementary Fig. 3). We note, however, that a comprehensive phosphoproteomics analysis of SARS-CoV-2-infected cells [57] suggests that other than interactions involving the N protein, which was found to be phosphorylated at multiple sites, most of the viral PPIs that were detected by Y2H analysis but not in our B2H system involve proteins that were not detectably phosphorylated.

### Genetic dissection of specific SARS-CoV-2 PPIs

A benefit of two-hybrid approaches for studying PPIs is that detected interactions can be readily dissected genetically, something that is particularly straightforward to do with our B2H system. As a proof-of-principle, we used a structure-based approach to investigate the effects of targeted mutations on specific SARS-CoV-2 PPIs, identifying substitutions that disrupt one interaction but not another. In addition to facilitating the evaluation of specific circulating or targeted mutations, our B2H system can readily be adapted to screen for randomly generated mutations that selectively affect one PPI and not another [58] when there is insufficient information to make informed predictions from structural or other data. The identification of such mutations could facilitate the functional analysis of particular PPIs and inform the choice of potential drug targets for small molecule drug design. Furthermore, with a suitably modified reporter strain to improve compound accessibility [59], compound or peptide libraries could be screened to identify candidates that might target specific SARS-CoV-2 PPIs. It should also be feasible to adapt our B2H reporter system for an *in vitro* cell-free protein expression system, thereby facilitating compound screenings.

### Use of oxidizing B2H reporter strain enables detection of RBD-ACE2 interaction

Based on previous studies, we anticipated a high-affinity interaction between the spike RBD and ACE2 [9,23,34]. With our modified bacteria-based system, we found that the RBD-ACE2 interaction resulted in a roughly 3-fold increase in *lacZ* reporter gene expression over background, a relatively modest effect. One possible explanation is that the λCI-ACE2 fusion protein is produced at relatively low levels compared with unfused λCI and other λCI fusion proteins we have studied in the past (Supplementary Fig. 8), perhaps resulting in intracellular concentrations insufficient to saturate the DNA-binding site on our *lacZ* reporter. Another possible explanation (not mutually exclusive) lies in the fact that both the SARS-CoV-2 spike protein and ACE2 are glycosylated in mammalian cells [9,60], with some studies suggesting that glycan-side chain interactions may be important in stabilizing the RBD-ACE2 interaction [61,62]. Thus, the interaction detected in our B2H system could be compromised by the lack of mammalian-like N- and O-glycosylation in *E. coli* [63].

Our B2H system enabled us to assess the effects of specific RBD amino acid substitutions that have been identified in globally circulating SARS-CoV-2 variants. We focused specifically on three VOCs, as designated by the CDC at the time we initiated our study: the Alpha variant (B.1.1.7), the Beta variant (B.1.351) and the Epsilon variant (B.1.429), carrying RBD substitutions N501Y, N501Y/K417N/E484K, and L452R, respectively (see Fig. 5D) [41]. We note that as of September 21, 2021 each of these variants has been deescalated from a VOC to a variant being monitored (VBM) by the CDC. We also note that the highly contagious and rapidly proliferating Delta variant, currently designated as a VOC, harbors the L452R substitution in the RBD, together with a second substitution (T478K) [64].

Mutated in both the Alpha and the Beta variants, residue N501 is localized at the binding interface with ACE2 [34,65] and many reports have suggested that the N501Y substitution increases the affinity of the RBD for ACE2 [23,48,66–70] (but see [71] for a discrepant prediction), potentially explaining the elevated infectivity of the Alpha variant. In a study in which the effects of all possible RBD amino acid substitutions were examined using a yeast-surface-display platform, Starr *et al.* identified N501Y as one of the substitutions causing the highest gain in ACE2-binding affinity [23]. In contrast, our B2H assay did not reveal any significant effect of the N501Y substitution on the strength of the RBD-ACE2 interaction. Possibly this discrepancy is due to the lack of glycosylation in the bacterial system; in fact, an ACE2 glycan (N322) that has been reported to enhance RBD-ACE2 binding is part of the same binding patch that includes N501 [62]. Nonetheless, we did observe a binding enhancing, compensatory effect of the N501Y substitution when tested in the context of the Beta variant. That is, we found that Beta-associated substitutions K417N and E484K both reduced ACE2 binding when tested individually, and that the N501Y substitution compensated for these effects, partially in the case of E484K and fully in the case of K417N. Our results thus suggest that substitutions K417N and E484K, which have been implicated in significant immune escape [43–45,72–74], may impose a cost on ACE2 binding that is compensated by the N501Y substitution (see also [70]). Consistent with our findings, the K417N substitution has been previously reported to weaken ACE2 binding [72,75,76]; however, in contrast with our results, Starr *et al.* [23] found that the E484K substitution had a small positive effect on ACE2 binding.

In the case of the L452R substitution, which is present in the Epsilon variant and also in the Delta variant [64], we observed a modest enhancement of ACE2 binding. Residue L452 is positioned at the edge of the binding interface with ACE2 and although this residue does not make direct contact with ACE2 [34,77], evidence suggests that substitution L452R enhances viral infectivity significantly [77,78]. Furthermore, it has been suggested that the L452R mutation is responsible for the dramatic clonal expansion of lineages carrying this mutation [79], possibly due to a decrease in the potency of antibody neutralization or through other immune escape characteristics [44,46,64,77,78,80]. Whether or not an effect of the L452R substitution on ACE2 binding, apparently modest, is a contributing factor in the rapid spread of variants carrying this mutation remains to be determined.

## Summary

Taken together, our results illustrate the utility of a B2H system as an accessible and economical genetic tool to complement other methods for studying viral PPIs. To the best of our knowledge, we provide the first bacteria-based viral interactome, describing sixteen different intraviral PPIs from SARS-CoV-2. As a non-eukaryotic system, the B2H assay is unlikely to contain bridging factors that can complicate the interpretation of positive results. At the same time, the bacterial system lacks the machinery for enabling potentially relevant post-translational modifications such as protein phosphorylation (which could however be engineered into the system; [53]) and protein glycosylation. Although generally a limitation, the lack of protein glycosylation could in certain situations be informative, enabling a comparison between systems that do and do not support this modification. The new oxidizing B2H reporter strain that we describe enabled us to detect the SARS-CoV-2 spike RBD-ACE2 interaction and characterize the effects of several RBD substitutions present in circulating variants. This strain provides a means to test newly arising coronavirus lineages for binding to ACE2 or other human cell surface receptors in the future, as well as extending the reach of the B2H system to include disulfide bond-dependent PPIs in general.

## Supplementary Figure Legends

**Supplementary Fig. 1: Fusion constructs of positive B2H interactions**. List of all B2H-identified SARS-CoV-2 PPIs. Green-shaded rectangles indicate the plasmid from which Protein A (first-mentioned protein in the left column) is produced to elicit a positive interaction with Protein B encoded by the other plasmid. When an interaction of Protein A and Protein B was identified regardless of the encoding plasmid, rectangles are shaded green in the column “both”.

**Supplementary Fig. 2: Medium-to-weak SARS-CoV-2 protein-protein interactions identified B2H assays**. Shown are two-hybrid data for medium-to-weak interactions (Miller unit values between 2- and 5-fold above the negative control Miller unit value). Indicated ORFs are fused either to the αNTD (indicated as α) or to full-length λCI (indicated as CI). α and λCI negative controls express full-length a and full-length λCI, respectively. The interaction of domain 4 of the RNAP σ^70^ subunit (fused to the αNTD) with the flap domain of the RNAP β subunit (fused to λCI) served as a positive control (pos) [81,82]. Bar graphs show the averages of three biological replicates (n=3) and β-galactosidase activities are given in Miller units. Error bars indicate the standard deviation. Values indicated with asterisks are significantly different from the negative control. ***: P<0.001; ****: P<0.0001 (One-way ANOVA with Turkey’s multiple comparison test).

**Supplementary Fig. 3: Comparison of B2H with Y2H and co-IP data.** Interaction matrix showing all SARS-CoV-2 PPIs identified with the B2H assay (this study) or with the Y2H and co-IP assays [6]. Squares designating PPIs identified solely by the B2H assay are colored purple and contain a white disc, whereas squares designating PPIs identified solely by the Y2H assay or by co-IP experiments are colored red and blue, respectively. Squares designating interactions that were identified by two or three of the assays contain the respective colors as indicated in the key at the left side of the matrix.

**Supplementary Fig. 4: Comparison of SARS-CoV-2 B2H data with SARS-CoV-1 two-hybrid data.** Interaction matrix showing the sixteen PPIs identified with the B2H assay (this study) and indicating which of these PPIs were also identified in at least one of three previous SARS-CoV-1 two hybrid studies, including two Y2H analyses [7,27] and one mammalian two-hybrid analysis [26]. Squares designating PPIs identified solely by the B2H assay are colored purple and contain a white disc. Coloring within the central disc designates a PPI that was also identified in one or two of the SARS-CoV-1 two-hybrid analyses (von Brunn *et al*. [7], orange; Imbert *et al*. [27], yellow; Pan *et al*. [26], cyan). Note that none of the PPIs were identified in all three SARS-CoV-1 analyses.

**Supplementary Fig. 5: Western blot analysis showing intracellular levels of Nsp10, Nsp14 and Nsp16 fusion proteins.** (top) Anti-αNTD and (bottom) anti-λCI western blot of cell lysates taken from over-night cultures of cells used for B2H analysis shown in **Fig. 4B**. Samples in lanes 1-5 are from cells producing the indicated λCI-Nsp10 fusion protein or λCI (see key) and the α-Nsp16 fusion protein, whereas samples in lanes 10-14 are from cells producing the indicated λCI-Nsp10 fusion protein or λCI and the α-Nsp14 fusion protein. Samples in lanes 6-9 are from cells producing the indicated λCI-Nsp10 fusion protein and full-length α. We note that substitutions D29A+S33A appeared to be mildly destabilizing, whereas substitutions F16A+F19A+V21A appeared to be mildly stabilizing.

**Supplementary Fig. 6: Western blot analysis showing intracellular levels of Nsp10, Nsp15 and Nsp16 fusion proteins.** (top) Anti-αNTD and (bottom) anti-λCI western blot of cell lysates taken from over-night cultures of cells used for the subsequent B2H analysis shown in **Fig. 4D**. Samples in lanes 1-5 are from cells producing the indicated α-Nsp16 fusion protein or α (see key) and the λCI-Nsp10 fusion protein, whereas samples in lanes 10-14 are from cells producing the indicated α-Nsp16 fusion protein or α and the λCI-Nsp15 fusion protein. Samples in lanes 6-9 are from cells producing the indicated α-Nsp16 fusion protein and full-length λCI. Lanes indicated with “X” are not discussed in the current manuscript but were not cut from the blot to avoid excessive manipulations of the original image. Note: Native, chromosomally encoded full-length α is detected in all samples.

**Supplementary Fig. 7: Domains of the SARS-CoV-2 spike protein**. Depiction of SARS-CoV-2 spike domains, including the signal peptide (SP, predicted from aa 1-13 or 1-16), the N-terminal domain (NTD, aa 1-330), the receptor binding domain (RBD, aa 331-521), the C-terminal domain (CTD, aa 522-1273) and the transmembrane domain (TMD, aa 1202-1273). The table lists the spike domains that were produced in *E. coli* (as B2H fusion proteins) and tested for interaction with ACE2, their precise corresponding loci on the SARS-CoV-2 genome, and the amino acids encoded by each test domain.

**Supplementary Fig. 8: Western blot analysis showing intracellular levels of spike RBD fusion proteins in oxidizing vs. reducing *E. coli* test strains.** (top) Anti-αNTD and (bottom) anti-λCI Western blot of cell lysates taken from over-night cultures of BLS148 or B2H cells used for the subsequent two-hybrid analyses shown in **Fig. 5B**. Samples in lanes 1-5 and 10-14 are from BLS148 cells and B2H cells, respectively, producing the indicated α-RBD fusion protein or α (see key) and the λCI-ACE2 fusion protein, whereas samples in lanes 6-9 and 15-18 are from BLS148 cells and B2H cells, respectively, producing the indicated α-RBD fusion protein and fulllength λCI. Lanes indicated with “X” are not discussed in the current manuscript but were not cut from the blot to avoid excessive manipulations of the original image. White dotted line indicates stitched image where extraneous material was removed. Note: Native, chromosomally encoded full-length α is detected in all samples.

**Supplementary Fig. 9: Western blot analysis showing intracellular levels of spike RBD mutants.** (top) Anti-α-NTD and (bottom) anti-λCI western blot of cell lysates taken from overnight cultures of BLS148 cells used for the subsequent two-hybrid analysis shown in **Fig. 5C**.

Samples in lanes 1-9 are from cells producing the indicated a-RBD fusion protein or a (see key) and the λCI-ACE2 fusion protein, whereas samples in lanes 10-17 are from cells producing the indicated a-RBD fusion protein and full-length λCI. Note: Native, chromosomally encoded fulllength a in all samples.

**Supplementary Table Legends**

**Supplementary Table 1:** List of (**A**) strains and plasmids, and (**B**) oligonucleotide primers used in this study.

## Acknowledgments

We thank Eleanor Fleming, Zoí Feder, Kemardo Henry, EmilyKate McDonough, Hanif Vahedian Movahed and Simon Dove for valuable discussion; Simon Dove and Jonathan Abraham for comments on the manuscript; and Sydney Rosa Teixeira for technical support. BLS, PD and AH were supported by a “Maximizing Investigators’ Research Award” (MIRA; No. R35GM136247) awarded to AH.

## Author contribution

AH and BLS designed the study. BLS performed the experimental work with support from PD. GG analyzed crystal structure data and provided predictions for the mutational screens. BLS and AH drafted the manuscript with contributions from all coauthors.

## Data availability

All data generated during and/or analyzed during the current study are either provided within the manuscript or are available from the corresponding authors upon reasonable request.

## Material and Methods

### Bacterial strains and growth conditions

*E. coli* strains MAX Efficiency™ DH5αF’IQ (Invitrogen) and NEB^®^ 5-alpha F’IQ (New England Biolabs, NEB) were used for routine cloning procedures and chemically competent *E. coli* were transformed with plasmid DNA by the standard heat shock procedure. FW102 O_L_2-62 and BLS148 strains were used for bacterial two-hybrid assays. All strains listed in Supplementary Table 1A were grown in LB medium containing the appropriate antibiotics at standard concentrations. BLS148 was created by P1 phage transduction of *ΔlacIZYA::kmR* from strain TB12 (P1 phage lysates were a gift from Thomas Bernhardt, Harvard Medical School) to SHuffle^®^ Express (NEB) according to a protocol established by Robert T. Sauer (Massachusetts Institute of Technology; protocol available at: https://openwetware.org/wiki/Sauer:P1vir_phage_transduction), generating BLS128. Deletion of *lacZ* in BLS128 was verified by colony PCR (using primers oBLS107+oBLS108 targeting *lacZ* to test for absence of *lacZ* and primers oBLS109+oBLS110 targeting *motA* and primers oBLS138+oBLS139 targeting *cyaA* as control reactions). Next, TSS competent BLS128 cells were created according to [83], transformed with pCP20 (encoding the yeast Flp recombinase gene to flip out the kanamycin resistance gene) and grown over night at 30 °C on LB plates containing carbenicillin (100 μg/ml; Carb100). The next day, 10 colonies were picked and restreaked on LB plates without antibiotics and then grown over night at 42 °C. From each of those strains a single colony was picked and re-streaked on LB plates containing either Carb100, kanamycin (20 μg/ml; Km20) or spectinomycin (50 μg/ml, Sp50) and grown over night at 30 °C. A single Sp50 resistant but Carb100 and Km20 sensitive colony was picked and reverified by streaking on the same growth plates. This strain that had lost the *kmR* resistance cassette was then designated BLS133. Finally, the β-galactosidase reporter present on the F’ was introduced into BLS133 by mating with strain FW102 O_L_2-62 [84]. For this, both BLS133 and FW102 O_L_2–62 were grown over night at 37 °C in LB Sp50 or Km20, respectively, and then streaked on top of each other on the same LB plate. After about 8 hours at 37 °C cells were resuspended in LB and plated in serial dilutions on LB plates containing Sp50, Km20 and X-gal (40 μg/ml; X-gal40) and then grown over night at 37 °C. A blue colony was picked and reverified by streaking again on a LB plate containing Sp50, Km20 and X-gal40, creating BLS148, a bacterial two-hybrid-compatible SHuffle^®^ Express strain.

### Plasmid construction

All plasmids generated in this study (see Supplementary Table 1A) were either constructed using standard restriction enzyme-based cloning procedures or by Gibson assembly. Gibson assembly was performed for 1 h at 50 °C by default. Primers employed for plasmid construction are listed in Supplementary Table 1B. Plasmid sequence integrity was verified by Sanger sequencing from Genewiz or Quintarabio (both Boston, MA, USA). Unless otherwise stated, all sequence templates, except for Nsp11, were ordered as *E. coli* codon-optimized gene fragments from Twist Bioscience (San Francisco, CA, USA).

Except for spike, Nsp2, Nsp3, RNA-Polymerase (Nsp12) and helicase (Nsp13), all full-length codon-optimized gene fragments were digested with NotI+BamHI, purified by DNA Clean & Concentrator kit (Zymo Research) and then ligated into 50 ng NotI+BamHI-digested pBRα or pACλCI using T4 ligase (NEB) according to standard protocols generating the plasmids listed in Supplementary Table 1A.

#### Spike

For pS63, the spike full-length (FL) sequence was amplified from *E. coli* codon-optimized gene fragments by SARS_67+SARS_68 and cloned into NotI+BamHI-digested pBRα by Gibson assembly. For pS64, the NTD sequence was amplified from pS63 by SARS_67+SARS_69 and then cloned into NotI+BamHI-digested pBRα by Gibson assembly. For pS65, the RBD sequence was amplified from pS63 by SARS_70+SARS_71 and then cloned into NotI+BamHI-digested pBRα by Gibson assembly. For pS66, the CTD sequence was amplified from pS63 by SARS_68+SARS_72 and then cloned into NotI+BamHI-digested pBRα by Gibson assembly. For pS67, the Ectodomain (aa 13-1213) was amplified from pS63 by SARS_73+SARS_74 and then cloned into NotI+BamHI-digested pBRα by Gibson assembly. For pS68, the Ectodomain (aa 16-1213) was amplified from pS63 by SARS_74+SARS_75 and then cloned into NotI+BamHI-digested pBRα by Gibson assembly. pS70 was generated by site-directed mutagenesis (SDM; see below) using primers SARS_17+SARS_76 and pS63 as a template. For pS72, FL spike was amplified from pS63 by SARS_77+SARS_78 and then cloned into NotI+BamHI-digested pACλCI by Gibson assembly. For pS73, the NTD sequence was amplified from pS63 by SARS_77+SARS_79 and then cloned into NotI+BamHI-digested pACλCI by Gibson assembly. For pS74, the RBD sequence was amplified from pS63 by SARS_80+SARS_81 and then cloned into NotI+BamHI-digested pACλCI by Gibson assembly. For pS75, the CTD sequence was amplified from pS63 by SARS_78+SARS-82 and then cloned into NotI+BamHI-digested pACλCI by Gibson assembly. For pS76, the Ectodomain (aa 131213) was amplified from pS63 by SARS_83+SARS_84 and then cloned into NotI+BamHI-digested pACλCI by Gibson assembly. For pS77, the Ectodomain (aa 16-1213) was amplified from pS63 by SARS_84+SARS_85 and then cloned into NotI+BamHI-digested pACλCI by Gibson assembly. pS79 was generated by SDM using primers SARS_17+SARS_86 and pS63 as a template.

#### Nsp2

The Nsp2 sequence was ordered as two single gene fragments, which were further amplified by PCR using primers SARS_109+SARS_110 or SARS_111+SARS_112 and then cloned into NotI+BamHI-digested pBRωGP by Gibson assembly, creating pS85. Next, the whole Nsp2 open reading frame (ORF) was cut from pS85 by NotI+BamHI and inserted into 50 ng NotI+BamHI-digested pBRa or pACλCI using T4 ligase (NEB) according to standard protocols, creating pS179 and pS180, respectively.

#### Nsp3

The Nsp3 sequence was ordered as four single gene fragments, which were further amplified by PCR using primers SARS_115+SARS_116, SARS_117+SARS_118, SARS_119+SARS_120 or SARS_121+SARS_122 and then cloned into NotI+BamHI-digested pBRωGP by Gibson assembly, creating pS89. Next, the whole Nsp3 open reading frame (ORF) was cut from pS89 by NotI+BamHI and inserted into 50 ng NotI+BamHI-digested pBRa or pACλCI using T4 ligase (NEB) according to standard protocols, creating pS181 and pS182, respectively.

#### RNA-Polymerase (Nsp12)

The RNA-Polymerase sequence was ordered as three single gene fragments, which were further amplified by PCR using primers SARS_131+SARS_132, SARS_133+SARS_134 or SARS_135+SARS_136 and then cloned into NotI+BamHI-digested pBRωGP by Gibson assembly, creating pS173. Next, the whole Nsp3 open reading frame (ORF) was cut from pS89 by NotI+BamHI and inserted into 50 ng NotI+BamHI-digested pBRa or pACλCI using T4 ligase (NEB) according to standard protocols, creating pS181 and pS182, respectively.

#### helicase (Nsp13)

The helicase sequence was ordered as two single gene fragments, which were further amplified by PCR using primers SARS_125+SARS_126 or SARS_127+SARS_128 and then cloned into NotI+BamHI-digested pBRωGP by Gibson assembly, creating pS169. Next, the whole Nsp2 open reading frame (ORF) was cut from pS85 by NotI+BamHI and inserted into 50 ng NotI+BamHI-digested pBRa or pACλCI using T4 ligase (NEB) according to standard protocols, creating pS221 and pS222, respectively.

#### ACE2

The ACE2 N-terminal peptidase domain (aa 19-615) was ordered as a single gene fragment and then further amplified by PCR using primers SARS_264+SARS_265 or SARS_266+SARS_267 and then cloned into 50 ng NotI+BamHI-digested pBRα or pACλCI by Gibson assembly, creating pS260 and pS261, respectively.

<u>Nsp11</u> was cloned into pBRα or pACλCI as annealed primers. For this, 10 μl of 100 μM SARS_139 and SARS_140 primers were mixed with 1 μl T4 Polynucleotide kinase (PNK; NEB) in 1x PNK reaction buffer (NEB). The reaction mix was placed in a BioRad T100 Thermal cycler, incubated for 30 min at 37 °C, inactivated for 5 min at 95 °C and then cooled to 4 °C at a 0.1 °C/s ramp rate. The annealed oligos were diluted 1:50 and then ligated into 50 ng NotI+BamHI-digested pBRα or pACλCI using T4 ligase (NEB), generating pS197 and pS198, respectively.

### Plasmid mutagenesis

Plasmid mutagenesis to create SARS-CoV-2 mutant genes was achieved using the Q5^®^ Site-Directed Mutagenesis (SDM) Kit according to the manufacturer’s instructions (NEB) or by using Gibson assembly with mutations introduced into the complementary overhang regions of the primer sequences. For the Gibson assembly, plasmids were amplified with the indicated primer pairs and 1 μl of the resulting PCR reaction was then ligated by Gibson assembly in a 10 μl reaction volume.

pS254 was generated by SDM using primers SARS_253+SARS_254 and plasmid pS215 as a template. pS256 was generated by SDM using primers SARS_256+SARS_257 and plasmid pS215 as a template. pS257 was generated by SDM using primers SARS_256+SARS_257 and plasmid pS254 as a template. pS262 was generated by Gibson assembly using primers SARS_268+SARS_269 and plasmid pS196 as a template. pS263 was generated by Gibson assembly using primers SARS_270+SARS_271 and plasmid pS196 as a template. pS264 was generated by Gibson assembly using primers SARS_272+SARS_273 and plasmid pS196 as a template. pS267 was generated by Gibson assembly using primers SARS_280+SARS_281 and plasmid pS65 as a template. pS271 was generated by Gibson assembly using primers SARS_287+SARS_288 and plasmid pS65 as a template. pS272 was generated by Gibson assembly using primers SARS_289+SARS_290 and plasmid pS65 as a template. pS273 was generated by Gibson assembly using primers SARS_289+SARS_290 and plasmid pS271 as a template. pS275 was generated by Gibson assembly using primers SARS_295+SARS_296 and plasmid pS65 as a template. pS276 was generated by Gibson assembly using primers SARS_291+SARS_292 and plasmid pS65 as a template. pS277 was generated by Gibson assembly using primers SARS_293+SARS_294 and plasmid pS65 as a template. pS278 was generated by Gibson assembly using primers SARS_291+SARS_292 and plasmid pS267 as a template. pS279 was generated by Gibson assembly using primers SARS_293+SARS_294 and plasmid pS267 as a template. pS280 was generated by Gibson assembly using primers SARS_293+SARS_294 and plasmid pS278 as a template.

### β-galactosidase assays

β-galactosidase assays to study the SARS-CoV-2 interactome were performed essentially as described previously [85]. In particular, pBRa and pACλCI plasmids containing the indicated inserts were co-transformed into FW102 OL2-62 by the heat shock procedure. Briefly, 2 μl of each plasmid (1:10 dil.) were mixed with 20 μl chemically competent FW102 O_L_2-62 cells, incubated on ice in 96-well PCR plates (VWR) for 30 min and then heat-shocked for 1 min at 42 °C in a BioRad T100 Thermal cycler. Cells were placed on ice for 5 min, recovered in 80 μl fresh LB medium and then incubated at 37 °C for 1 h (Please note: we found that commercially available premixed LB drastically reduces transformation efficiency and also subsequent over night culture growth; we thus recommend using non-premixed LB medium instead). The 96-well plates were sealed with Rayon Films (VWR) to allow proper aeration and prevent contamination. Afterwards, 50 μl of transformed cells were transferred to 2 ml deep well plates containing 500 μl LB Carb100, chloramphenicol (25 μg/ml; Cm25), Km20 and 5 μM IPTG and grown over night at 37 °C, 800 rpm. The next day, 4 μl over-night culture was transferred to 96-well flat bottom microtiter plates containing 200 μl LB Carb100, Cm25, Km20 and 20 μM IPTG and grown until approx. OD_600_ 0.15-0.2 (measured in a VERSA Max microplate reader, Molecule Devices, San Jose, CA, USA). Then, 20 μl lysis solution (for one 96-well plate mix: 1.2 ml PopCulture^®^ Reagent (MilliporeSigma, MA, USA), 2.5 μl 400 U/μl rLysozyme™ (MilliporeSigma, MA, USA) and 1.25 μl Benzonase^®^ Nuclease (MilliporeSigma, MA, USA)) was added to the cells and incubated for at least 30 min at 37 °C and 800 rpm (longer incubation times were found to not negatively affect the experimental results). Afterwards, 30 μl lysed cell suspension was added to a fresh 96-well flat-bottom microtiter plate containing 150 μl Z-buffer/ONPG solution (60 mM Na_2_HPO_4_, 40 mM NaH_2_PO_4_, 10 mM KCl, 1 mM MgSO_4_, 1 mg/ml ortho-Nitrophenyl-β-galactoside (ONPG)) and OD_420_ values were recorded in a VERSA Max microplate reader (Molecule Devices, San Jose, CA, USA). β-galactosidase activity in Miller units was then calculated as described previously [85].

β-galactosidase assays to study the RBD-ACE2 interaction were performed as follows. Strain BLS148 was transformed with the appropriate plasmids, as described in the preceding paragraph. Upon recovery of the transformed cells for 1 h at 37 °C, 50 μl cells were then transferred to 500 μl LB Carb100, Km20, Cm25 and 50 μM IPTG and grown for approx. 20 h at 30 °C and 800 rpm. Subsequently, 15 μl cells were transferred to 185 μl LB medium in 96-well microtiter plates, combined with 20 μl lysis solution (for one 96-well plate: 1.2 ml PopCulture^®^ Reagent (MilliporeSigma, MA, USA), 5.0 μl 400U/μl rLysozyme™ (MilliporeSigma, MA, USA) and 2.5 μl Benzonase^®^ Nuclease (MilliporeSigma, MA, USA)) and incubated for at least 30 min at 30 °C, 800 rpm. All subsequent steps were then performed as described above.

### Western blot analysis

To verify the production of the respective fusion proteins, western blots of cell lysates from over-night cultures were performed. For this, co-transformed cells were grown in the indicated IPTG concentration over-night in 550 μl total volume in 2 ml deep well plates at 30 or 37 °C, 800 rpm. The next day, OD_600_ values were recorded and 500 μl cells were pelleted by centrifugation (1 min, 21,000 x *g*, room temperature (RT)) and either stored at −80 °C or directly processed. Cell pellets were then resuspended in lysis buffer (BugBuster^®^ Protein Extraction Reagent (MilliporeSigma, MA, USA) supplemented with 1x cOmplete™, EDTA-free Protease Inhibitor Cocktail (MilliporeSigma, MA, USA), 1 U/μl rLysozyme™ (MilliporeSigma, MA, USA; final concentration) and 0.5 U/μl Benzonase^®^ Nuclease (MilliporeSigma, MA, USA, final concentration)). The amount of lysis buffer for each cell pellet was calculated as follows: μl lysis buffer = OD_600_ x ml of culture pelleted x 60. Cells were lysed for 30 min at RT in an overhead shaker. Next, lysed cells were mixed 1:5 in PBS (10.14 mM Na_2_HPO_4_, 1.76 mM NaH_2_PO_4_, 2.7 mM KCl, 137 mM NaCl; pH 7.4, Boston Bioproducts, MA, USA) and then incubated in 1 x Laemmli SDS sample buffer (Boston Bioproducts, MA, USA) at 95 °C for 10 min. 10 μl of the resulting solution was then applied to either 4-12% Criterion™ XT Bis-Tris Protein Gels (BioRad, Hercules, CA, USA) or NuPAGE™ 4 – 12% Bis-Tris Mini Protein Gels (Thermo Fisher Scientific, MA, USA). Upon gel separation, proteins were transferred to Amersham Protran 0.45 NC nitrocellulose membranes (Cytiva, MA, USA) using a Trans-Blot Turbo Transfer System (BioRad Hercules, CA, USA), blocked in blocking buffer (TBST: 50 mM Tris-HCl, 150 mM NaCl, pH 7.4, 0.1 % Tween-20 supplemented with 5% non-fat dry milk) for 30 min at RT and then incubated with mouse anti-a-NTD and rabbit anti-CI primary antibodies (both 1:3,000 dil.) in blocking buffer for 1 h at RT. After washing with TBST, blots were incubated with IRDye^®^ 680RD goat anti-mouse and IRDye^®^ 800CW goat anti-rabbit (both 1:10,000 dil.; LI-COR Biosciences, NE, USA) in blocking buffer for 1 h at RT in the dark. After washing with TBST, proteins were then detected using a ChemiDoc MP system (BioRad. Hercules, CA, USA).

### Protein crystal structure analysis

Interfaces of two protein complexes, SARS-CoV-2 Nsp16-Nsp10 (PDB ID: 6W4H [31]) and SARS-CoV-1 Nsp10-Nsp14 (PDB ID: 5NFY [29]), were analyzed using PDBePISA software *(insert ref 20).* Amino acids involved in hydrogen bond formation or substantially contributing to hydrophobic contacts in each complex were subjected to alanine mutagenesis and tested in B2H assays. Structural images were prepared using PyMOL software (Schrodinger, LLC. 2010. The PyMOL Molecular Graphics System, Version 2.4.1).

### Statistical analysis

Presentation of bacterial two-hybrid data and statistical analysis using one-way or two-way ANOVA with Tukey’s or Dunnett’s multiple comparison test was done using GraphPad Prism (v. 9.1.2; San Diego, CA, USA).

